# Persistent cross-species SARS-CoV-2 variant infectivity predicted via comparative molecular dynamics simulation

**DOI:** 10.1101/2022.04.18.488629

**Authors:** Madhusudan Rajendran, Gregory A. Babbitt

## Abstract

Widespread human transmission of SARS-CoV-2 highlights the substantial public health, economic, and societal consequences of virus spillover from wildlife and also presents a repeated risk of reverse spillovers back to naïve wildlife populations. We employ comparative statistical analyses of a large set of short-term molecular dynamic (MD) simulations to investigate potential human to bat (*Rhinolophus macrotis*) cross-species infectivity allowed by the binding of SARS-CoV-2 receptor-binding domain (RBD) to angiotensin-converting enzyme 2 (ACE2) across the bat progenitor strain and emerging human strain variants of concern (VOC). We statistically compare the dampening of atom motion during binding across protein sites upon the formation of the RBD/ACE2 binding interface using bat vs. human target receptors (i.e. bACE2 and hACE2). We report that while the bat progenitor viral strain RaTG13 shows some pre-adaption to binding hACE2, it also exhibits stronger overall affinity to bACE2. However, while the early emergent human strains and later VOC’s exhibit robust binding to both hACE2 and bACE2, the delta and omicron variants exhibit evolutionary adaption of binding to hACE2. However, we conclude there is a still significant risk of mammalian cross-species infectivity of human VOC’s during upcoming waves of infection as COVID-19 transitions from a pandemic to endemic status.

## Introduction

Coronavirus disease (COVID-19) is caused by novel severe acute respiratory syndrome coronavirus 2 (SARS-CoV-2). The virus first emerged from the Wuhan province in China [1,2]. Since its emergence, the virus has had a devastating effect on the world’s population, resulting in more than 5.5 million deaths worldwide [3]. Since being declared a global pandemic by the World Health Organization (WHO) on March 11, 2020, the virus continues to cause devastation, with several countries enduring multiple waves of outbreaks of this viral illness. A variety of mathematical model types, including statistical, deterministic, stochastics and agent-based models have used to study the transmission dynamics and control of COVID-19 [4–7].Traditional theory applied to the viral epidemiology of COVID-19 have extended the compartmental modeling approach pioneered by Kermack and McKendrick (i.e. SIR model) [8] as well as the concept of the evolutionary arms race derived from evolutionary game theory introduced by Maynard-Smith [9] as an extension Nash [10]. Modern advances in high-throughput DNA sequencing have allowed COVID-19 forecasting to be better parameterized with regards to real-time sequence-based surveillance as well as temporal-spatial patterns of behavior in viral-infected human populations [11]. Recently, the epidemiological modeling community has also recognized the need to model beyond simple viral infection rates in human populations and to incorporate information regarding human interactions with other species and environments that lead to zoonotic spillover events (e.g. the OneHealth framework) [12]. In this regard, some key questions of concern raised by the ongoing pandemic that are particularly difficult to address with compartmental modeling in epidemiology are (A) what are the general molecular properties of proteins that facilitate viral spillovers between two or more species?, (B) how does the random occurrence (i.e. neutral evolution) of these properties relate to the frequency or likelihood of zoonotic spillovers of viruses between species?, and (C) how long might it take after a spillover to humans for a highly virulent emergent strain to evolve to lose these properties and become a more benign endemic species specific strain?

The initial outbreak of COVID-19 was initially linked to a local seafood market in Wuhan, China, where the sale of wild animals has been implicated as the primary source of SARS-CoV-2 infections [13]. Furthermore, the current SARS-CoV-2 virus is known to have 96.2% similarity to the bat coronavirus RaTG13 at the whole genome level [14,15]. Based on the viral genome sequence and further evolutionary analysis, Chinese horseshoe bats of genus *Rhinolophus* have been pinpointed as the most likely natural reservoir host for the recent emergence of the SARS-CoV-2 virus [16]. However, as of now, no definitive intermediate host that may be more closely associated with humans (e.g. domestic pets or livestock) has been identified. Interestingly based on the isolation of closely related genomes from Malayan pangolins (Manis javanica), they are thought to be possible intermediate hosts of the SARS-CoV-2 via wildlife markets in Wuhan [17].

Since its initial spillover from *Rhinolophus* bats and subsequent introduction to the global human population, the genome of SARS-Cov-2 has mutated. As a result, thousands of variants of SARS-CoV-2 have emerged [18]. The WHO defined the SARS-CoV-2 variants of concern (VOC) as a variant with increased transmissibility, virulence, and decreased response to available diagnostics, vaccines, and therapeutics [19]. Based on the recent epidemiological updated by WHO, as of January 10, 2022, five SARS-CoV-2 VOCs have been identified since the pandemic’s beginning [20]. Alpha (B.1.1.7) was the first VOC described in the United Kingdom in late December 2020. Then came the beta (B.1.351) and Gamma (P.1) variants which were first reported in South Africa in December 2020 and Brazil in January 2021, respectively. Until most recently, Delta (B.1.617.2) has been the most dominant variant. It was first reported in India in December 2020. Lastly, the Omicron (B.1.1.529) variant has become the most dominant variant, with its origin in South Africa in late November 2021 [19]. \

The virus’s origin from the spillover of a zoonotic pathogen, and the broad host range of the virus is partly because its ACE2 target receptor is found in all major vertebrate groups [21]. The ubiquity of ACE2 coupled with the high prevalence of SARS-CoV-2 in the global human population explains the multiple spillback infections since the emergence of the virus in 2019. In spillback infection, the human hosts transmit the SARS-CoV-2 virus to cause infection in nonhuman animals. In addition to threating wildlife and domestic animals, the repeated spillback infection may lead to the establishment of new animal hosts from which SARS-CoV-2 can then pose a risk of secondary spillover infection to humans through bridge hosts or new established enzootic reservoirs. There is a small number of reports of human to animal transmission of SARS-CoV-2 in pet cats and dogs and gorillas, tigers, lions, and other felines in zoos in the USA, Europe, and South Africa [22,23]. Initial human to animal transmission has resulted in sustained outbreaks in farmed mink in Europe and North America, with likely mink to human transmission reported in the Netherlands [24,25]. More recently, there have been some accounts of reverse spillover of VOC strains, including omicron, into wild North American white-tailed deer populations [26–28]. Experimental infection of Egyptian fruit bats (Rousettus aegyptiacus) resulted in transient subclinical infection with oral and fecal shedding [29]. Given that the probable sources of SARS-CoV-2 or its progenitor are bat species, the potential risk of reverse zoonotic transmission from humans to bats and the subsequent negative impacts have to be recognized and studied. Furthermore, a significant concern in such secondary spillover events is the subsequent evolution of mutant strains leading to increased transmissibility and/or mortality in humans, reduced sensitivity to neutralizing antibodies, and reduced vaccine efficacy.

A very crucial and unresolved key question of concern regarding the continued evolution of this pandemic is whether and how long the human VOC’s remain capable of reverse spillovers into other species of mammals as they adapt their binding to more specifically target human ACE2. We have recently introduced new statistical applications for comparing the divergence of short-term rapid MD of proteins in functionally relevant molecular binding states (i.e. comparing the divergence of atom fluctuation between bound vs. unbound protein states) [30–32] and applied this to study of the evolution of emergent and endemic viral strains related to SARS-CoV-2 [33]. We have also recently applied this method to the study of the evolution of antibody binding escape mutations as well [34]. Here, we utilize this same comparative molecular dynamics-based approach to study of individual amino acid sites involved in the binding of the various strains of the SARS-CoV-2 viral receptor-binding domain (RBD) to both the human and *Rhinolophus* bat ACE2 orthologs (hACE2 and bACE2 resp.). We present multiple-test corrected site-wise statistical comparisons of the SARS-CoV-2 RBD binding signatures of all currently reported VOCs in the presence of both hACE2 and bACE2, identifying sites that have led to significantly increased hACE2 binding of SARS-CoV-2 related viral strains as they evolved during the course of the pandemic. We report that while some specific adaptations to hACE2 have emerged in the delta and omicron VOC, there is still a persistent risk of reverse spillover to bats and likely many other mammals even two years into the current pandemic.

## Materials and Methods

### PDB structure and model preparation

Structures of the RBD of spike protein, hACE2, and Greater Horseshoe Bat Rhinolophus macrotis bACE2 were obtained from the Protein Data Bank (PDB). The summary of the structures used for MD simulations is listed in Table 1. Upon downloading the structures from PDB, any crystallographic reflections and other small molecules used in crystallization were removed. Furthermore, any glycans present in the structure were also removed. During the cleanup of the PDB, any glycans were removed and then later rebuilt using glycoprotein builder [35] so that the PDB file structure regarding atom types was compatible with the Amber 20 preprocessing software tLeAP [36]. See details below. When preparing the structures, we needed each of the variants (RaTG13, Wuhan WT, Alpha, Beta, Delta, Kappa, Epsilon, and Omicron BA.1 and Omicron BA.2) bound to hACE2 and bACE2. We were able to find structures in the PDB where the RaTG13 variant was bound to bACE2 and the human variants bound to hACE2. Therefore, to model the RaTG13 variant bound to hACE2 and the human variants bound to bACE2, we used UCSF Chimera’s MatchMaker superposition tool to properly place the receptor belonging to the opposite species [37]. Using pdb4amber (AmberTools20), hydrogen atoms were added, and crystallographic waters were removed [36]. Any mossing loop structures in the files were inferred via homology modeling using the “refine loop” command to modeler within UCSF chimera [38,39]. The structure of omicron BA.2 RBD is not available on PDB, and therefore Alphafold2 was used to create a 3D structure, details of which are detailed below.

**Table 1:** Table summarizing the primary models (RBD variants bound to hACE2/bACE2) used for MD simulations. bACE2 denotes ACE2 of bat origin, and hACE2 denotes ACE2 of human origin.

| RBD Variant | PDB ID | Receptor | PDB ID |
| --- | --- | --- | --- |
| RaTG13 (BatCoV) | 7CN4 | bACE2 | 7C8J |
| RaTG13 (BatCoV) | 7CN4 | hACE2 | 6VW1 |
| Wuhan WT (nCoV) | 7C8D | bACE2 | 7C8J |
| Wuhan WT (nCoV) | 7C8D | hACE2 | 6VW1 |
| Alpha (B.1.1.7) | 7LWV | bACE2 | 7C8J |
| Alpha (B.1.1.7) | 7LWV | hACE2 | 6VW1 |
| Beta (B.1.351) | 7LYO | bACE2 | 7C8J |
| Beta (B.1.351) | 7LYO | hACE2 | 6VW1 |
| Delta (B.1.617.2) | 7V7Q | bACE2 | 7C8J |
| Delta (B.1.617.2) | 7V7Q | hACE2 | 6VW1 |
| Kappa (B.1.617.1) | 7V7E | bACE2 | 7C8J |
| Kappa (B.1.617.1) | 7V7E | hACE2 | 6VW1 |
| Epsilon (B.1.429) | 7N8H | bACE2 | 7C8J |
| Epsilon (B.1.429) | 7N8H | hACE2 | 6VW1 |
| Omicron (BA.1) | 7T9J | bACE2 | 7C8J |
| Omicron (BA.2) | 7T9J | hACE2 | 6VW1 |
| Omicron (BA.2) | Alphafold2 | bACE2 | 7C8J |
| Omicron (BA.2) | Alphafold2 | hACE2 | 6VW1 |

### Model glycosylation

As mentioned previously, glycans present in the original PDB structure were deleted. Predicted glycosylation was rebuilt for the Amber forcefield using the glycoprotein builder on the glycam.org web server [35]. The glycans were rebuilt using the GLYCAM-06j-1 force field [40]. 2-acetamido-2-deoxy-beta-D-glucopyranose was attached to ASN10 in the RBD of all variants. Similarly, 2-acetamido-2-deoxy-beta-D-glucopyranose was attached to ASN227, ASN264, and ASN503, and 2-acetamido-2-deoxy-beta-D-glucopyranose-(1-4)-2-acetamido-2-deoxy-beta-D-glucopyranose was attached to ASN720 of bACE2. Lastly, 2-acetamido-2-deoxy-beta-D-glucopyranose was attached to ASN271, and 2-acetamido-2-deoxy-beta-D-glucopyranose-(1-4)-2-acetamido-2-deoxy-beta-D-glucopyranose was attached to ASN221, ASN258, ASN490, and ASN714 of hACE2.

### Molecular dynamic simulation protocols

MD simulation protocol was followed as previously described, with slight modifications [30–32,41]. Briefly, for each MD comparison, large replicate sets of accelerated MD simulations were prepared and then conducted using the particle mesh Ewald method implemented on the graphical processor unit (GPU) hardware by pmemd.cuda (Amber20) [42–44]. The MD simulations were either performed on a Linux mint 19 operating system (two Nvidia RTX 2080 Ti or two Nvidia RTX 3080 Ti) or on high performance computing cluster (Nvidia A100). All comparative MD analysis via our DROIDS pipeline was based upon 100 replicate sets of 1 nanosecond accelerated MD run (i.e., 100×1ns MD run in each comparative state RBD, e.g., RBD bound to the receptor, unbound RBD). Explicitly solvated protein systems were first prepared using teLeap (AmberTools 20) using the ff14SB protein forcefield, in conjunction with the GLYCAM_06j-1 force field [40,45]. Solvation was generated using the Tip3p water model in a 12nm octahedral water box [46]. Automated charge neutralization was also done with teLeap software with Na+ and Cl-ions. Each replicate set was preceded by energy minimization, 300 picoseconds of heating to 300K, a ten nanosecond of equilibration, followed by random equilibration intervals for each replicate ranging from 0-0.5 nanoseconds. All simulations were regulated using the Anderson thermostat at 300K and one atmospheric pressure [47]. Root mean square atom fluctuations were calculated in CPPTRAJ using the atomicfluct command [48].

### Comparative protein dynamic analyses with DROIDS 4.0 and statistical analyses

Comparative signatures of dampened atom fluctuation during RBD binding to ACE2 were presented as protein site-wise divergence in atom fluctuation in the ACE2 bound versus unbound states for each RBD. Divergences were calculated using the signed symmetric Kullback-Leibler (KL) divergence calculation in DROIDS 4.0. Significance tests and p-values for these site-wise differences were calculated in DROIDS 4.0 using two-sample Kolmogorov-Smirnov tests with the Benjamini-Hochberg multiple test correction in DROIDS 4.0. The mathematical details of DROIDS 4.0 site-wise comparative protein dynamics analysis were published previously by our group and can be found here [30–32]. This code is available at our GitHub web landing: https://gbabbitt.github.io/DROIDS-4.0-comparativeprotein-dynamics/, which is also available at our GitHub repository https://github.com/gbabbitt/DROIDS-4.0-comparative-protein-dynamics.

### Omicron BA.2 receptor binding domain prediction

Using the derived RBD amino acid sequence of Omicron BA.2 (Accession # UJE45220.1), we used Alphafold2 to create a predicted protein structure. AlphaFold2 is a neural networkbased deep learning model which first searches for homologous sequences with existing structures to use as a scaffold on which to place the new sequence [49]. The AlphaFold2-based prediction was run with the “single sequence” mode using the predicted TM-score (PTM) method. We also specified that the algorithm should run an Amber relaxation procedure to repair any structural violations in the predicted model [50].

## Results

### Stronger binding of RatG13 RBD to bACE2 than to hACE2

We performed MD simulations of RaTG13 RBD bound and unbound to bACE2 and RaTG13 RBD bound and unbound to hACE2. To compare atomic fluctuations between RatG13 RBD bound and unbound structures, we used site-wise KL divergence along with multiple test corrected two-sample KS tests. The more negative the KL divergence value of a specific amino acid residue, the stronger the dampening of atomic fluctuations due to the RBD interactions with bACE2/hACE2. As one would expect, the bat RaTG13 RBD has a better binding with stronger amino acid-specific interactions with the bACE2 (Figure 1A, 1C). However, in the case of hACE2, dampening of atomic fluctuations is lesser at those specific sites due to ACE2 being of human origin (Figure 1A, 1D). Interestingly, the amino acid residues involved with the interactions of both bACE2 and hACE2 are very similar. This was observed in the normalized KL divergence graph, where the normalized KL divergence values per amino acid are similar between bACE2 and hACE2 (Figure 2A). Other studies have found that 26 residues of the bACE2 and nine residues of the RaTG13 RBD are present at the interface. These residues create 12 H-bonds, two salt bridges, and 157 non-bonded contacts [51]. The residues of RatG13 RBD that are strongly dampened by bACE2 include K417, L455, F456, S477, N487, and D501 (Figure 1A). The weaker dampening of atomic fluctuations of RaTG13 RBD and hACE2 is primarily due to the lesser number of interactions between the RBD and hACE2. Compared to bACE2, hACE2 only makes 113 non-bonded contacts and 9 H-bonds [52]. Lastly, MD comparison of RaTG13 bound to bACE2 and RaTG13 bound to hACE2 show that almost half of the amino acids in the RBD of RaTG13 behave statistically different (Figure 1B). When comparing RaTG13 RBD bound to hACE2 with RaTG13 RBD bound to bACE2, the significance tests are conducted site-wise. Therefore, a separate test was conducted at each given amino acid site to compare the significant difference in fluctuations of the backbone atoms. Thus, a separate D value from a two-sample KS test is obtained for each amino acid site, and a multiple test correction (Benjamini-Hochberg) was applied to adjust the p-value to account for multiple significance tests. Interestingly, there is no statistical difference in the atomic fluctuation dampening of some of the residues that interact with ACE2. In contrast, a majority of the interacting residues have statistically different atomic fluctuations between bACE2 and hACE2 (p < 0.001) (Figure 1B).

**Figure 1:**
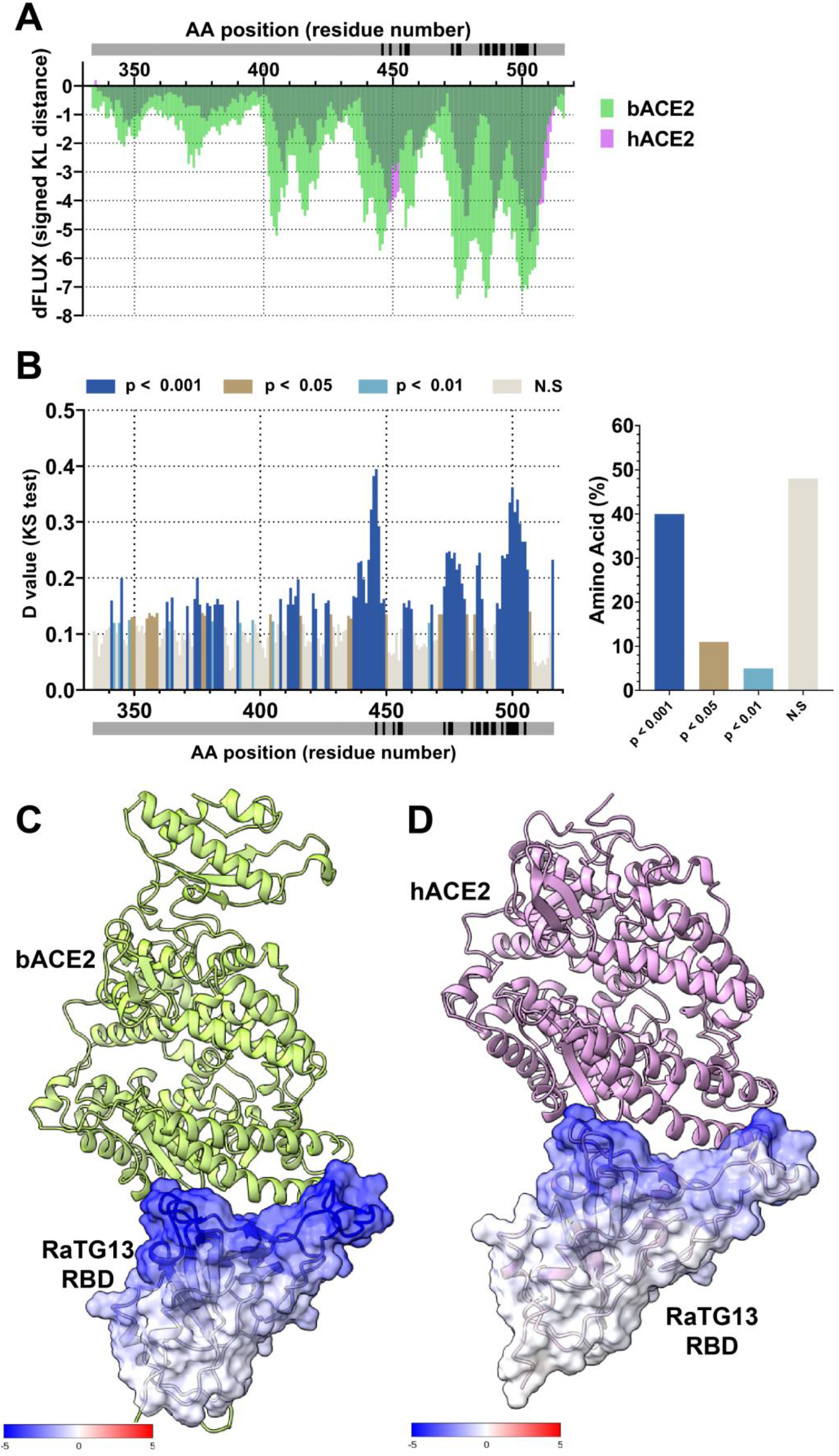
Analysis of atomic fluctuation differences of RaTG13 RBD bound to bACE2 and hACE2. (A) Sequence positional plotting of dampening of atom motion on RaTG13 RBD by bat ACE2 (bACE2, green) and human ACE2 (hACE2, pink). (B, left panel) Multiple test corrected two-sample KS tests of significance for the difference in atomic fluctuations of RaTG13 RBD bound to bACE2 and RaTG13 RBD bound to hACE2. The grey bar in (A) and (B) denotes the RBD domain amino acid backbone with the location of RBD residues interacting with ACE2 shown in black. (B, right panel) Percent of amino acid of the RaTG13 RBD with different levels of significance. N.S. denotes no significance. The change in atom fluctuation is due to the (C) bACE2 and (D) hACE2 interactions with RaTG13 RBD (PDB 7CN4). Dark blue denotes a KL divergence value of −5, with red denoting a KL divergence value of +5. bACE2 (PDB 7C8J) is shown in green, and hACE2 PDB 6VW1) shown in pink.

**Figure 2:**
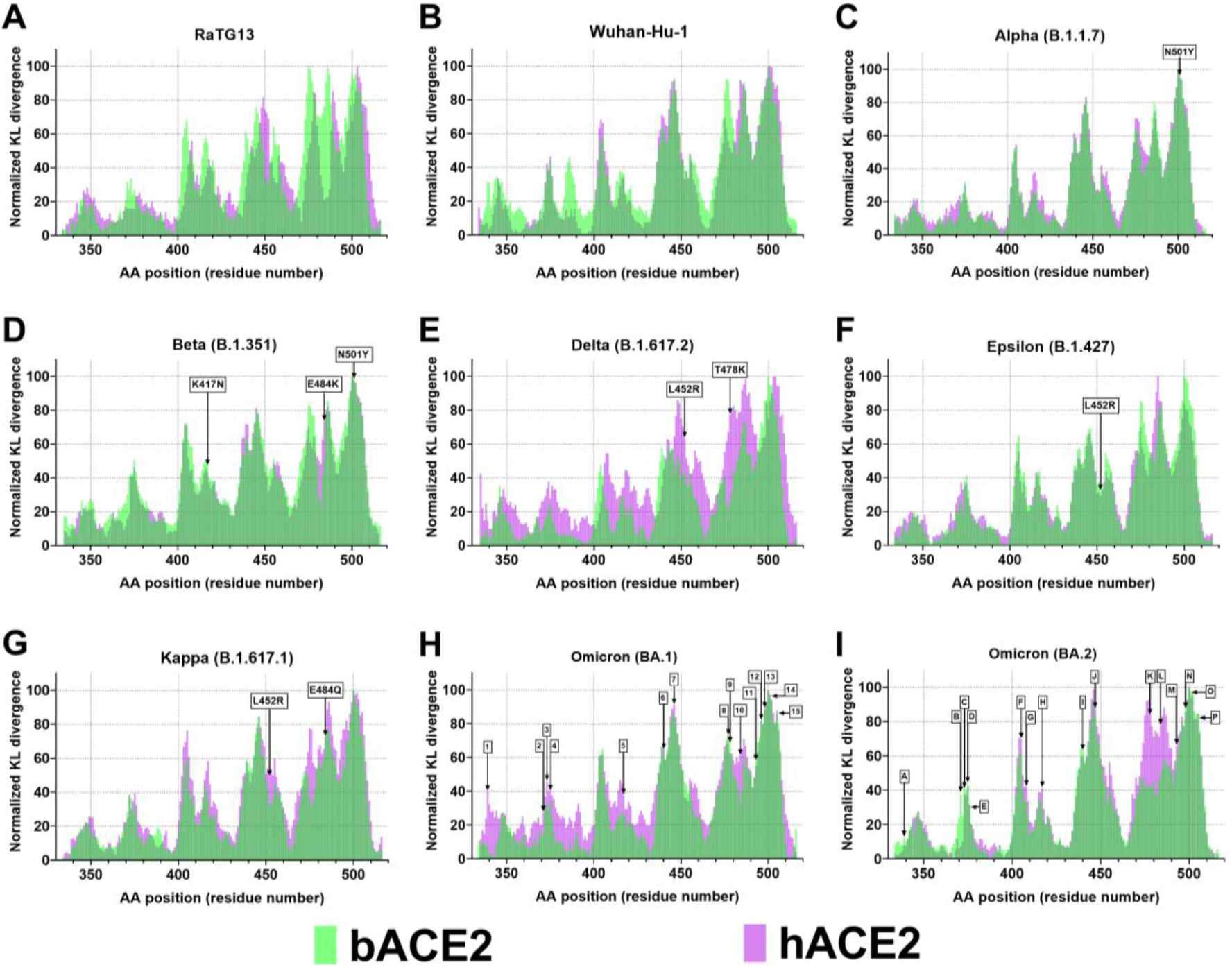
Binding interaction of the different SARS-CoV-2 variants with bACE2 and hACE2. Sequence positional plotting of normalized dampening of atom motion on (A) RaTG13, (B) Wuhan-Hu-1, (C) Alpha (B.1.1.7), (D) Beta (B.1.351), (E) Delta (B.1.617.2), (F) Epsilon (B.1.427), (G) Kappa (B.1.617.1), (H) Omicron (BA.1), and (I) Omicron (BA.2) RBDs by bat ACE2 (bACE2, green) and human ACE2 (hACE2, pink). The amino mutations denoted by the arrows correspond to the variant mutation. The list of omicron BA.1 mutations in (H) include (1) G339D, (2) S371L, (3) S373P, (4) S375F, (5) K417N, (6) N440K, (7) G446S, (8) S477N, (9) T478K, (10) E484A, (11) Q493K, (12) G496S, (13) Q498R, (14) N501Y, (15) Y505H. The list of omicron BA.2 in (I) include (A) G339D, (B) S371F, (C) S373P, (D) S375F, (E) T376A, (F) D405N, (G) R408S, (H) K417N, (I) N440K, (J) S447N, (K) T478K, (L) E484A, (M) Q493R, (N) Q498R, (O) N501Y, (P) Y505H.

### Compared with VBM, VOCs have better binding to hACE2 than to bACE2

In addition to RaTG13 RBD bound and unbound to bACE2 and hACE2, we performed MD simulations of other variants. These variants include the original Wuhan-Hu-1 strain, alpha (B.1.1.7), beta (B.1.351), delta (B.1.617.2), epsilon (B.1.427), kappa (B.1.1617.1), omicron (BA.1, omicron (BA.2). Similar to RaTG13, the MD simulation of the different variants included the RBD bound and unbound to bACE2 and hACE2. Upon calculating the site-wise KL divergence for the RBD bound/unbound of the different variants, we normalized the KL divergence values. The residue with very little atomic fluctuation dampening was set to 0, and the residue with the strongest atomic fluctuation dampening was set to 100 (Figure 2). Both RaTG13 and the original Wuhan-Hu-1 RBDs have a similar binding dynamic between the bACE2 and hACE2. The peaks of normalized KL divergence values between the bACE2 and hACE2 of the two variants being identical corresponds to the similar amino acid residues interacting with the bACE2 and hACE2 (Figure 2A, 2B). We performed a similar analysis with the variants being monitored (VBM). The VBM includes beta, epsilon, and kappa variants. Like the original bat progenitor and the first human variant, the VBM show identical normalized KL divergence plots between the hACE2 and bACE2. Even at the sites corresponding to the VBM mutation, the KL divergence values appear to be very similar (Figure 2D, 2F, 2G). Lastly, we also looked at the VOC’s interactions with bACE2 and hACE2. The VOC included alpha, delta, omicron BA.1 and omicron BA.2. Surprisingly, with the alpha variant, we see very similar normalized KL divergence plots between the bACE2 and hACE2 (Figure 2C). However, we see quite a difference in the delta and omicron variant plots. The normalized KL divergence values show additional peaks in the simulations with the hACE2. These additional peaks correspond to delta and omicron mutations (Figure 2E, 2H and 2I).

Lastly, to quantify the interaction of the different variants with bACE2 and hACE2, we also calculated the area under the curve (AUC) of the non-normalized KL divergence values, true atomic fluctuation dampening due to the RBD interaction with ACE2. As expected, the bat progenitor strain, RaTG13, has a higher AUC with bACE2 than hACE2. Since the Wuhuna-Hu-1 RBD is the first variant to make the transmission from bats to humans, the AUC profile is similar for both bACE2 and hACE2 (Figure 3A). Interestingly, all VOC (alpha, delta, omicron BA.1 and omicron BA.2) have a higher AUC for the hACE2 than bACE2. When a paired t-test was performed to compare the AUC values of the VOC bound to bACE2 and hACE2, we saw a significantly higher AUC profile of the VOC with hACE2 (Figure 3A, 3B). And lastly, there is no clear trend for the AUC profile of the VBM. This is also clearly seen with no significant difference in AUC values of VBM between bACE2 and hACE2 (Figure 3A, 3C).

**Figure 3:**
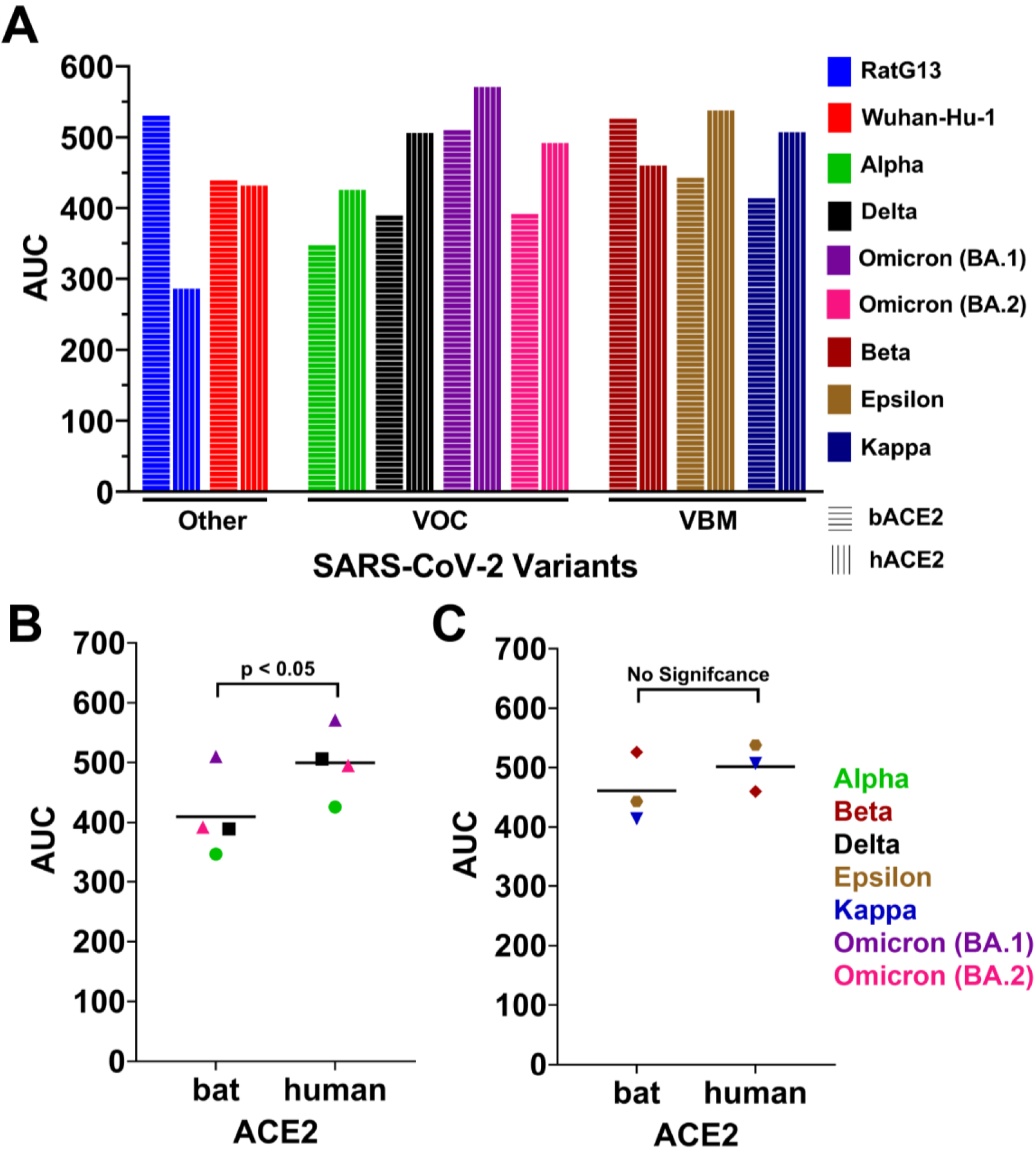
Binding dynamic differences between VOC and VBM with ACE2. (A) Area under the curve (AUC) values of the non-normalized KL divergence values of the different SARS-CoV-2 RBD bound and unbound to bACE2 and hACE2. Alpha, delta, and omicron variants are classified as variants of concern (VOC); beta, epsilon, and kappa variants are classified as variants being monitored (VBM). RaTG13 and Wuhan-Hu-1 are classified as others. Data points in (B)VOC and (C)VBM correspond to the AUC value of the different variants bound bACE2 and hACE2. The horizontal line corresponds to the mean value. A paired t-test was conducted to calculate the statistical significance.

### Stronger binding of Delta RBD to hACE2 than to bACE2

The delta variant (B.1.617.2) was first identified in India in October 2020. The two mutations that make up the RBD of the delta variant include L452R and T478K. The site-wise KL divergence interaction of the delta RBD between bACE2 and hACE2, denotes that hACE2 certainly has better binding dynamics than the bACE2 (Figure 4A). KS test between the delta RBD bound to bACE2 and delta RBD bound to hACE2 shows very few amino acids with statistically different atomic fluctuations (35%). About 65% of the amino acid of delta RBD behave very similarly when either bound to bACE2 or hACE2. At the sites corresponding to the delta variant mutation (L452R and T478K), we see a high level of significance in the difference in atomic fluctuation between the bACE2 and hACE2 (Figure 4B). As mentioned previously, when comparing delta RBD bound to hACE2 with delta RBD bound to bACE2, significance tests were conducted in a site-wise manner, with a D value from a two-sample KS test calculated for each amino acid, and Benjamini-Hochberg multiple test correction was applied to adjust the p-values. The test correction was done to account for the multiple significance tests. Lastly, compared to bACE2, the binding of the hACE2 also has a dampening effect on the residues that are farther away from the RBD/ACE2 interface (Figure 4C and 4D).

**Figure 4:**
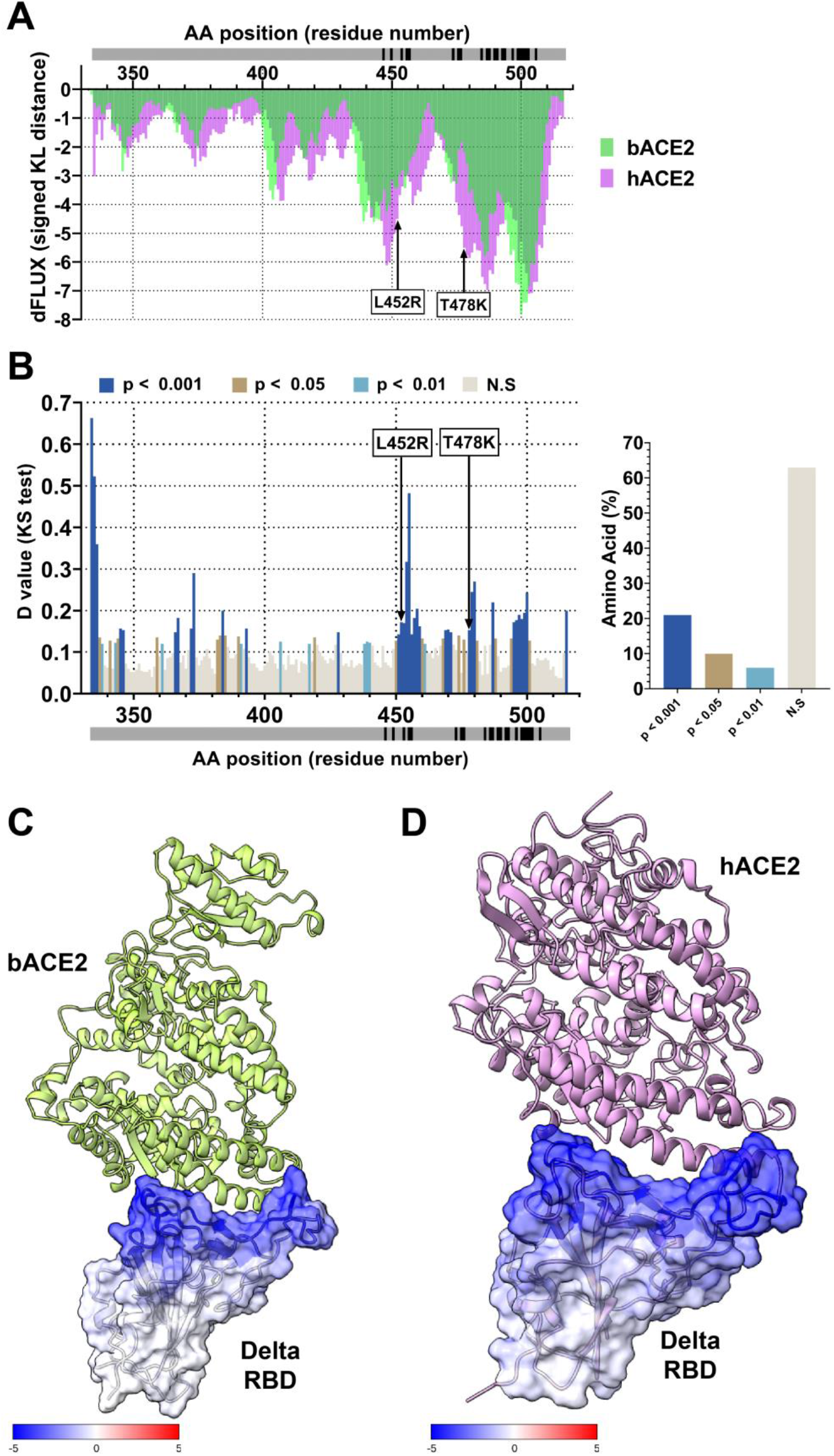
Analysis of atomic fluctuation differences of Delta (B.1.617.2) RBD bound to bACE2 and hACE2. (A) Sequence positional plotting of dampening of atom motion on delta RBD by bat ACE2 (bACE2, green) and human ACE2 (hACE2, pink). (B, left panel) Multiple test corrected two-sample KS tests of significance for the difference in atomic fluctuations of delta RBD bound to bACE2 and RaTG13 RBD bound to hACE2. The grey bar in (A) and (B) denotes the RBD domain amino acid backbone with RBD residues of interaction with ACE2 shown in black. (B, right panel) Percent of amino acid of the delta RBD with different levels of significance. N.S. denotes no significance. Arrows in (A) and (B) correspond to the delta variant mutations (L452R and T478K). The change in atom fluctuation is due to the (C) bACE2 and (D) hACE2 interactions with delta RBD (PDB 7V7Q). Dark blue denotes a KL divergence value of −5, with red denoting a KL divergence value of +5. bACE2 (PDB 7C8J) is shown in green, and hACE2 (PDB 6VW1) is pink.

### Omicron RBD mutations influence binding to hACE2

Unlike the delta variant, the omicron variant was most recently identified in South Africa in November 2021. Compared to all other variants, omicron variants include atleast 15 mutations in the RBD. Comparison of the atomic fluctuation of the omicron BA.1 RBD bound to bACE2 and hACE2 revealed certain amino acids with stronger dampening of atomic fluctuations when bound to hACE2 (Figure 5A). Furthermore, statistically significant differences in the atomic fluctuation of the omicron BA.1 RBD bound to bACE2 and hACE2 are only observed in 30% of the amino acids of the omicron. Of those different amino acid positions, 9 of them correspond to the omicron BA.1 RBD mutations (G339D, S371L, S373P, S345F, K417N, E484A, Q493K, G496S, and Q498R) (Figure 5B). Similar to the delta variant, the RBD of the omicron variant BA.1 was furthermore stabilized when bound to hACE2. This can be seen by the dampening of residues by hACE2 that are farther away from the RBD/hACE2 interface (Figure 5D). However, in the case of the bACE2, there is only dampening happening at the RBD/bACE2 interface (Figure 5C). Identical analysis of the more recent omicron BA.2 variant shows even more binding efficacy (Figure 6 A-D). The binding signatures of human and bat orthologs of ACE2 in response the alpha, delta and omicron viral RBD are also given in Supplemental Figure 1 and 2 and reflect similar trends of enhanced binding during the evolution of the binding interface.

**Figure 5:**
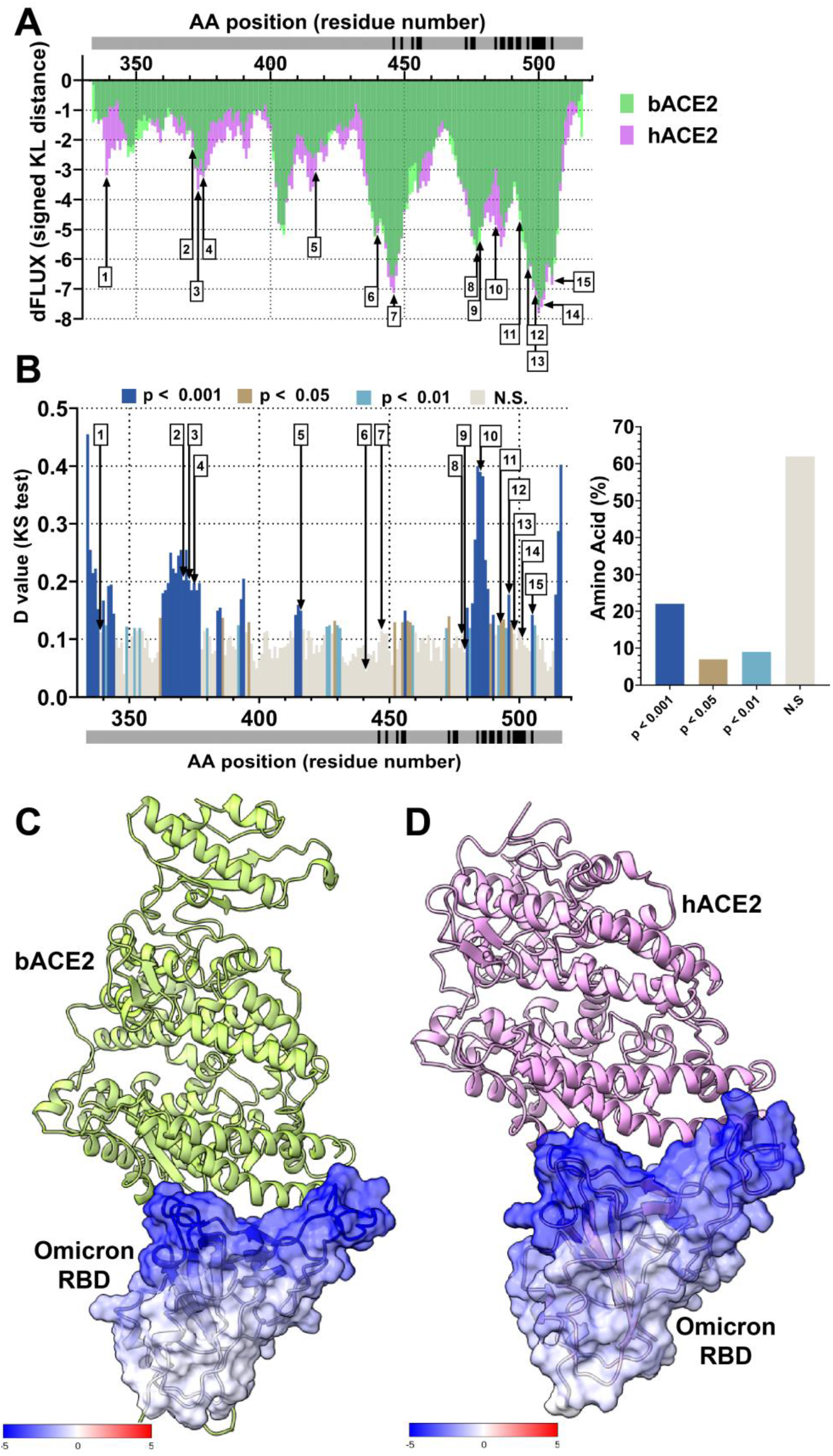
Analysis of atomic fluctuation differences of Omicron BA.1 variant RBD bound to bACE2 and hACE2. (A) Sequence positional plotting of dampening of atom motion on omicron RBD by bat ACE2 (bACE2, green) and human ACE2 (hACE2, pink). (B, left panel) Multiple test corrected two-sample KS tests of significance for the difference in atomic fluctuations of omicron RBD bound to bACE2 and omicron RBD bound to hACE2. The grey bar in (A) and (B) denotes the RBD domain amino acid backbone with RBD residues of interaction with ACE2 shown in black. (B, right panel) Percent of amino acid of the omicron RBD with different levels of significance. N.S. denotes no significance. Arrows in (A) and (B) correspond to the omicron variant mutations. The list of omicron mutations includes (1) G339D, (2) S371L, (3) S373P, (4) S375F, (5) K417N, (6) N440K, (7) G446S, (8) S477N, (9) T478K, (10) E484A, (11) Q493K, (12) G496S, (13) Q498R, (14) N501Y, (15) Y505H. The change in atom fluctuation is due to the (C) bACE2 and (D) hACE2 interactions with omicron RBD (PDB 7T9K). Dark blue denotes a KL divergence value of −5, with red denoting a KL divergence value of +5. bACE2 (PDB 7C8J) is shown in green, and hACE2 (PDB 6VW1) shown in pink.

**Figure 6:**
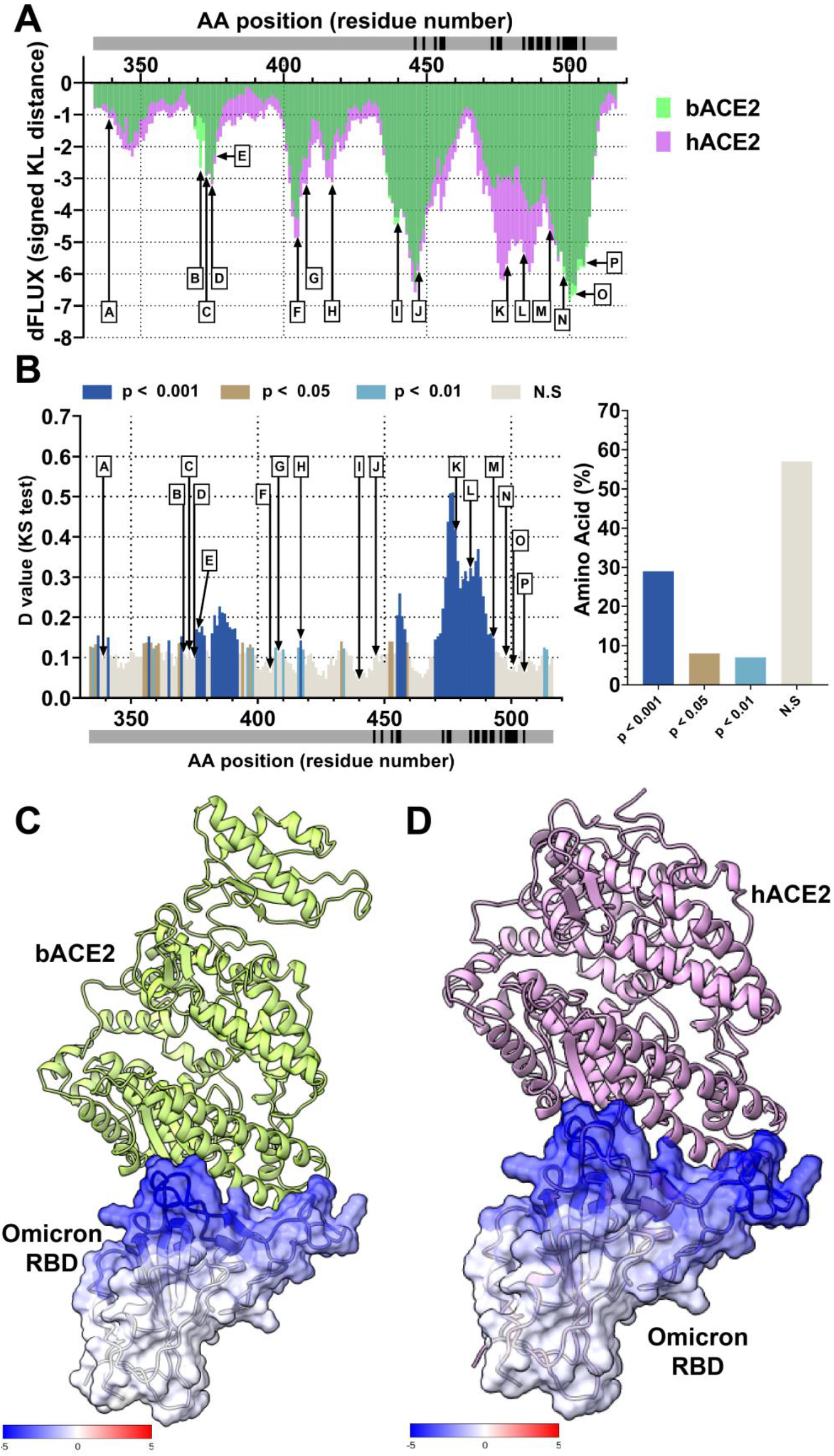
Analysis of atomic fluctuation differences of Omicron BA.2 variant RBD bound to bACE2 and hACE2. (A) Sequence positional plotting of dampening of atom motion on omicron RBD by bat ACE2 (bACE2, green) and human ACE2 (hACE2, pink). (B, left panel) Multiple test corrected two-sample KS tests of significance for the difference in atomic fluctuations of omicron RBD bound to bACE2 and omicron RBD bound to hACE2. The grey bar in (A) and (B) denotes the RBD domain amino acid backbone with RBD residues of interaction with ACE2 shown in black. (B, right panel) Percent of amino acid of the omicron RBD with different levels of significance. N.S. denotes no significance. Arrows in (A) and (B) correspond to the omicron variant mutations. The list of omicron mutations includes (A) G339D, (B) S371F, (C) S373P, (D) S375F, (E) T376A, (F) D405N, (G) R408S, (H) K417N, (I) N440K, (J) S447N, (K) T478K, (L) E484A, (M) Q493R, (N) Q498R, (O) N501Y, (P) Y505H. The change in atom fluctuation is due to the (C) bACE2 and (D) hACE2 interactions with omicron RBD (PDB 7T9K). Dark blue denotes a KL divergence value of −5, with red denoting a KL divergence value of +5. bACE2 (PDB 7C8J) is shown in green, and hACE2 (PDB 6VW1) shown in pink.

## Discussion / Conclusion

Using our comparative MD simulation pipeline, we compared the binding profile of bACE2 and hACE2 with the RBD domains of different SARS-CoV-2 variants (RaTG13, Wuhan-Hu-1, alpha, beta, delta, kappa, epsilon, omicron BA.1 and omicron BA.2). MD simulations of RatG13 spike protein revealed a better binding profile and stronger dampening of amino acids on the interface of RBD and bACE2 compared to hACE2. In the case of the RBD from Wuhan-Hu-1, the original 2019 SARS-CoV-2 virus, we see no difference in the binding profile between bACE2 and hACE2. Lastly, we also observed two different profiles with the VBM and VOC. In the case of VBM (beta, kappa, and epsilon variants), we see a similar binding profile and atomic fluctuation dampening in the RBD/bACE2 and RBD/hACE2 interfaces. On the other hand, VOCs (alpha, delta, and omicron variants) show preferable binding to hACE2 than to bACE2, indicating higher AUC values when RBD is bound to hACE2 than RBD bound to bACE2. The SARS-CoV-2 virus was first reported from pneumonia patients of the Wuhan city in China’s Hubei province. The spillover of SARS-CoV-2 from animals to humans occurred at the beginning of December 2019, when some of the pneumonia patients were involved in the wet animal market in the Hunnan district [53]. Genomic sequences, homology of ACE2 receptor, and single intact ORF on gene 8 of the virus indicate bats as the natural reservoir of these viruses. However, an unknown animal is yet to be identified as an intermediate host [54,55]. It should be noted that even though the initial spread of the disease was due to a spillover even, the rapid spread of the disease was primarily due to human-to-human transmission [53].

The receptor usage by the corona virus has been well known to be a significant determinant of host range, tissue tropism, and pathogenesis. Therefore, it is reasonable to assume that SARS-CoV-2 can infect humans, bats, and other species. As a matter of fact, several in vivo infection and seroconversion studies have confirmed that SARS-CoV-2 can infect rhesus monkeys, feline, ferret, and canines [56,57]. Our MD simulation has shown that RaTG13 RBD can bind to both bACE2 and hACE2, however to bACE2 with stronger binding, shown by severe dampening of atomic fluctuations in the bACE2/RBD interface (Figure 1A). Even though hACE2 doesn’t severely dampen the atomic fluctuations, the KL divergence profile is very similar to that of bACE2, indicating the flexibility of the virus to jump hosts. Additionally, it has also been shown that the RatG13 RBD Y493 is speculated to confer a potential steric clash to hACE2 [58].

As mentioned previously, for zoonotic spillover events to occur, humans must be exposed to the viruses. This can occur through direct contact with viruses excreted from infected bats or bridge hosts or through other contacts with infected animals such as slaughtering or butchering. The nature and intensity of the bat–human interface are critical to determining spillover risk. Human behavior is a primary determinant of exposure, which may increase contact with bats or with other animals (bridge hosts) that may expose susceptible humans. Little is known about the specific conditions of coronavirus spillovers. Still, human behaviors that may increase viral exposure include activities such as bat hunting and consumption, guano farming, and wildlife trading [59–61] (51-53). Due to recent spillover events, we see an identical binding profile with both bACE2 and hACE2 in the case of Wuhan-Hu-1 RBD (see Figure 2B). In the case of Wuhan-Hu-1 RBD, our MD simulation shows identical dampening of the atomic fluctuations and that the amino acid backbone in the RBD interacts with both bACE2 and hACE2 identically.

Regardless, in the case of human SARS-CoV-2 strains, we see a slightly different trend. In the case of VOC, including alpha, delta, and omicron variants, we see a slightly higher binding profile with hACE2 than bACE2 (Figure 3A, 3B, 4–6). However, unlike RaTG13 RBD, we do not see very strong differences in the atomic fluctuations in the bACE2/hACE2 interface with the VOC RBDs. We don’t see any significant differences in the binding profile with the VBM, including beta, epsilon and kappa variants. At some areas of the RBD, the atomic fluctuations of the VBM are very similar between bACE2 and hACE2. The ability of the recent VOC and VBM to bind to both hACE2 and bACE2 with only slight differences supports the reverse zoonosis theory. The transmission of SARS-CoV-2 from humans to numerous animals and conducted in vitro infection experiments make it clear that the virus can infect and be transmitted between a wide range of distantly related mammal species. For example, case reports on cats (Felis catus) living in the same household with COVID-19 patients in Europe, Asia, North America, and South America revealed that these animals could be infected with SARS-CoV-2, showing clinical manifestations ranging from asymptomatic to severe respiratory illness [62,63] (54, 55). The reports show that 14% of tested cats in Hong Kong were SARS-CoV-2 positive by RT-PCR.[64]

Furthermore, the seroprevalence screening performed among pets living in SARS-CoV-2-positive households in Italy demonstrated that 3.3% of dogs and 5.8% of cats were seropositive [65]. The high seroprevalence and SARS-CoV-2 detection rates in cats and, to some extent, in dogs indicate that these animals can be infected with SARS-CoV-2 [66]. Several other zoo animals, like tigers, lions, cougars, and gorillas, were found to test positive for the virus. Farmed minks are highly susceptible to SARS-CoV-2 infection, and, in some cases, they have transmitted the virus back to humans. SARS-CoV-2-positive minks were detected in 290 fur farms in Denmark, 69 mink fur farms in the Netherlands, 13 of 40 mink farms in Sweden, 23 out of 91 mink farms in Greece, 17 fur farms in the USA, four farms in Lithuania, two farms in Canada, and one fur farm in Italy, Latvia, Poland, France, and Spain [67,68]. As a result of the virus being able to infect multiple species and is also able to jump hosts, there are concerns that the introduction and circulation of new virus strains in humans could result in modifications of transmissibility or virulence and decreased treatment and vaccine efficacy.

In conclusion, our MD simulations identified that the original bat progenitor RaTG13 RBD shows preferential binding to its host bACE2 receptor than hACE2. However, some of the recent human variants show differential binding between bACE2 and hACE2. Lastly, the VOC RBD show slightly higher binding to hACE2 than the bACE2. These findings provide evidence that recent SARS-CoV-2 variants may infect bats and that the extensive species diversity of bats may have profound effects on SARS-CoV-2 evolution. Overall, our results indicate the possible inclusion of MD-based surveillance of the virus and the viral receptor, in addition to genomic surveillance.

## Acknowledgements

We would like to acknowledge Research Computing at Rochester Institute of Technology for providing computational resources and support.

---

**Supplemental Figure 1.**
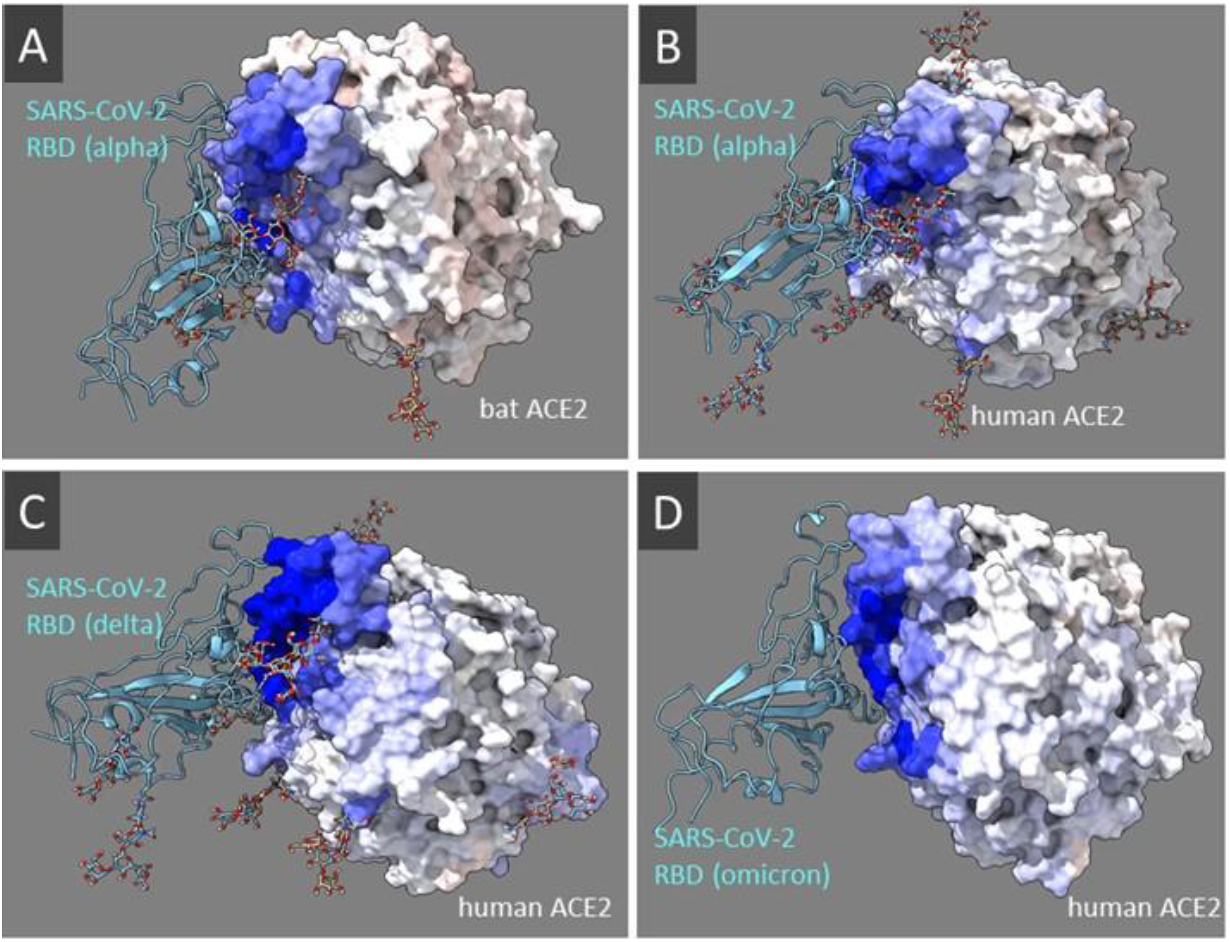
The binding signatures of SARS-CoV-2 receptor binding domain respective to human and bat (Rhinolophus macrotis) ACE2 orthologs. The blue mapping indicates dampened atom fluctuation calculated as the signed KL divergence between site-wise distributions of atom fluctuation in the viral-bound vs unbound state of ACE2. The bindingsignatures for (A) reverse spilloverof the SARS-CoV-2 alpha strain to bat ACE2 is compared to binding of (B) alpha, (C) delta, and (D) omicron strains to human ACE2.

**Supplemental Figure 2.**
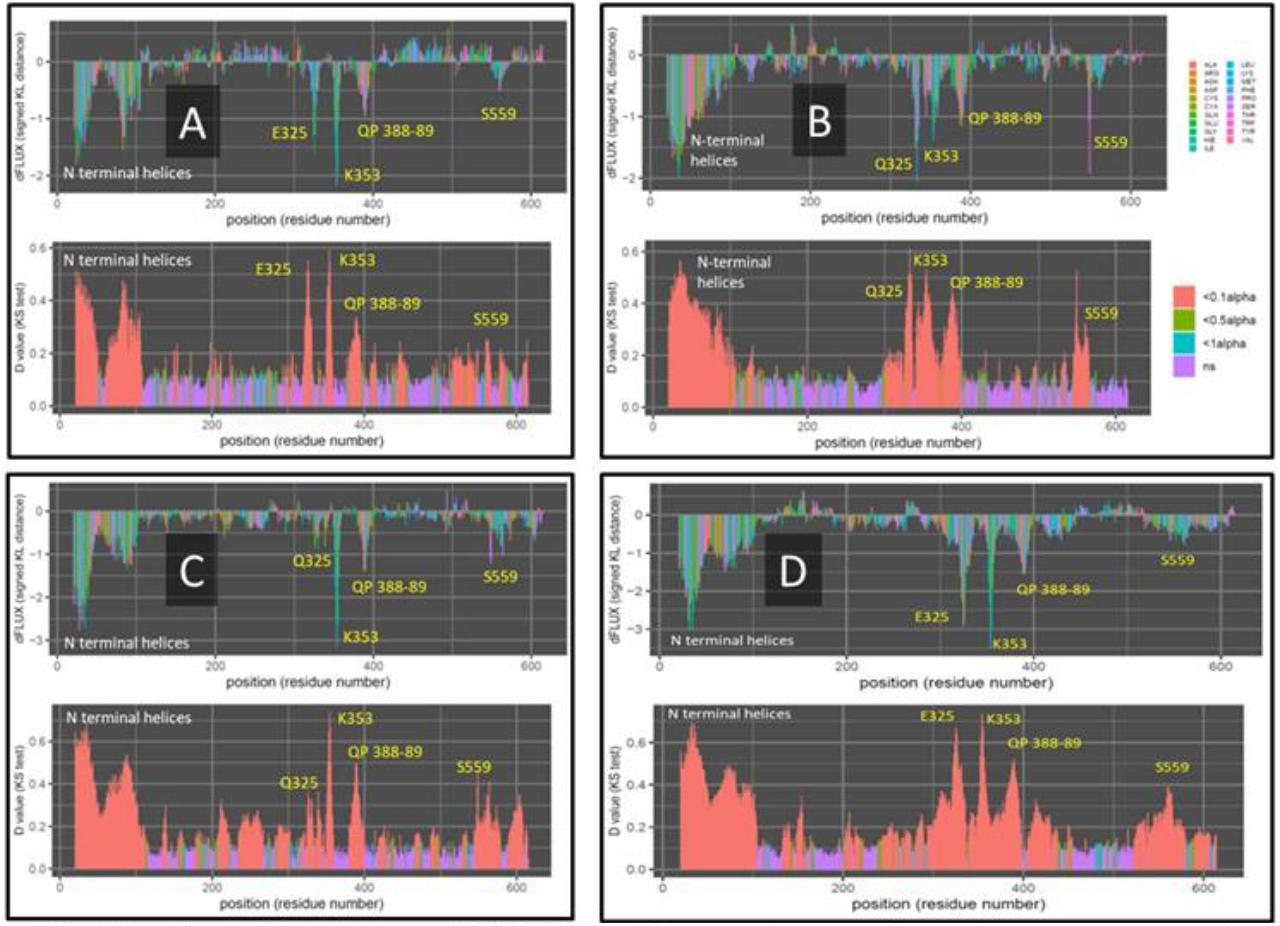
The binding signatures of SARS-CoV-2 receptor binding domain respective to human and bat (Rhinolophus macrotis) ACE2 orthologs. The top plot indicates dampened atom fluctuation calculated as the signed KL divergence between site-wise distributions of atom fluctuation in the viral-bound vs unbound state of ACE2. The bottom plot indicates where these differences in atom fluctuation are significantly different accordingto Benjamini-Hochberg correscted two-sample Kolmogorov-Smirnovtests. The bindingsignatures for (A) reverse spilloverof the SARS-CoV-2 alpha strain to bat ACE2 is compared to bindingof (B) alpha, (C) delta, and (D) omicron strains to human ACE2.

## Notes

### Competing Interest Statement

The authors have declared no competing interest.

## References

1. Huang C et al. 2020 Clinical features of patients infected with 2019 novel coronavirus in Wuhan, China. The Lancet 395, 497–506. (doi:10.1016/S0140-6736(20)30183-5)

2. Li Q et al. 2020 Early Transmission Dynamics in Wuhan, China, of Novel Coronavirus–Infected Pneumonia. N. Engl. J. Med. 382, 1199–1207. (doi:10.1056/NEJMoa2001316)

3. In press. WHO Coronavirus (COVID-19) Dashboard. See https://covid19.who.int (accessed on 26 March 2022).

4. Hellewell J et al. 2020 Feasibility of controlling COVID-19 outbreaks by isolation of cases and contacts. Lancet Glob. Health 8, e488–e496. (doi:10.1016/S2214-109X(20)30074-7)

5. Srivastava A, Chowell G. 2020 Understanding Spatial Heterogeneity of COVID-19 Pandemic Using Shape Analysis of Growth Rate Curves. MedRxiv Prepr. Serv. Health Sci., 2020.05.25.20112433. (doi:10.1101/2020.05.25.20112433)

6. Xue L, Jing S, Miller JC, Sun W, Li H, Estrada-Franco JG, Hyman JM, Zhu H. 2020 A data-driven network model for the emerging COVID-19 epidemics in Wuhan, Toronto and Italy. Math. Biosci. 326, 108391. (doi:10.1016/j.mbs.2020.108391)

7. Eikenberry SE, Mancuso M, Iboi E, Phan T, Eikenberry K, Kuang Y, Kostelich E, Gumel AB. 2020 To mask or not to mask: Modeling the potential for face mask use by the general public to curtail the COVID-19 pandemic. Infect. Dis. Model. 5, 293–308. (doi:10.1016/j.idm.2020.04.001)

8. Kermack WO, McKendrick AG, Walker GT. 1927 A contribution to the mathematical theory of epidemics. Proc. R. Soc. Lond. Ser. Contain. Pap. Math. Phys. Character 115, 700–721. (doi:10.1098/rspa.1927.0118)

9. Maynard Smith J. 1974 The theory of games and the evolution of animal conflicts. J. Theor. Biol. 47, 209–221. (doi:10.1016/0022-5193(74)90110-6)

10. Nash JF. 1950 Equilibrium points in n-person games. Proc. Natl. Acad. Sci. 36, 48–49. (doi:10.1073/pnas.36.1.48)

11. Rahimi I, Chen F, Gandomi AH. 2021 A review on COVID-19 forecasting models. Neural Comput. Appl., 1–11. (doi:10.1007/s00521-020-05626-8)

12. Kleczkowski A, Hoyle A, McMenemy P. 2019 One model to rule them all? Modelling approaches across OneHealth for human, animal and plant epidemics. Philos. Trans. R. Soc. B Biol. Sci. 374, 20180255. (doi:10.1098/rstb.2018.0255)

13. Lake MA. 2020 What we know so far: COVID-19 current clinical knowledge and research. Clin. Med. Lond. Engl. 20, 124–127. (doi:10.7861/clinmed.2019-coron)

14. Zhou P et al. 2020 A pneumonia outbreak associated with a new coronavirus of probable bat origin. Nature 579, 270–273. (doi:10.1038/s41586-020-2012-7)

15. Zhou P et al. 2020 Addendum: A pneumonia outbreak associated with a new coronavirus of probable bat origin. Nature 588, E6. (doi:10.1038/s41586-020-2951-z)

16. Zhang T, Wu Q, Zhang Z. 2020 Probable Pangolin Origin of SARS-CoV-2 Associated with the COVID-19 Outbreak. Curr. Biol. 30, 1346–1351.e2. (doi:10.1016/j.cub.2020.03.022)

17. Xiao K et al. 2020 Isolation of SARS-CoV-2-related coronavirus from Malayan pangolins. Nature 583, 286–289. (doi:10.1038/s41586-020-2313-x)

18. Biswas SK, Mudi SR. 2020 Genetic variation in SARS-CoV-2 may explain variable severity of COVID-19.Med. Hypotheses 143, 109877. (doi:10.1016/j.mehy.2020.109877)

19. Choi JY, Smith DM. 2021 SARS-CoV-2 Variants of Concern. Yonsei Med. J. 62, 961–968. (doi:10.3349/ymj.2021.62.11.961)

20. In press. Tracking SARS-CoV-2 variants. See https://www.who.int/health-topics/typhoid/tracking-SARS-CoV-2-variants (accessed on 26 March 2022).

21. Chou C-F et al. 2006 ACE2 orthologues in non-mammalian vertebrates (Danio, Gallus, Fugu, Tetraodon and Xenopus). Gene 377, 46–55. (doi:10.1016/j.gene.2006.03.010)

22. Leroy EM, Ar Gouilh M, Brugère-Picoux J. 2020 The risk of SARS-CoV-2 transmission to pets and other wild and domestic animals strongly mandates a one-health strategy to control the COVID-19 pandemic. One Health 10, 100133. (doi:10.1016/j.onehlt.2020.100133)

23. McAloose D et al. 2020 From People to Panthera: Natural SARS-CoV-2 Infection in Tigers and Lions at the Bronx Zoo. mBio 11, e02220–20. (doi:10.1128/mBio.02220-20)

24. Oreshkova N et al. 2020 SARS-CoV-2 infection in farmed minks, the Netherlands, April and May 2020. Euro Surveill. Bull. Eur. Sur Mal. Transm. Eur. Commun. Dis. Bull. 25. (doi:10.2807/1560-7917.ES.2020.25.23.2001005)

25. Oude Munnink BB et al. 2021 Transmission of SARS-CoV-2 on mink farms between humans and mink and back to humans. Science 371, 172–177. (doi:10.1126/science.abe5901)

26. Palmer MV et al. 2021 Susceptibility of white-tailed deer (Odocoileus virginianus) to SARS-CoV-2. J. Virol., JVI.00083–21. (doi:10.1128/JVI.00083-21)

27. Chandler JC et al. 2021 SARS-CoV-2 exposure in wild white-tailed deer (Odocoileus virginianus). Proc. Natl. Acad. Sci. 118, e2114828118. (doi:10.1073/pnas.2114828118)

28. Hale VL et al. 2022 SARS-CoV-2 infection in free-ranging white-tailed deer. Nature 602, 481–486. (doi:10.1038/s41586-021-04353-x)

29. Schlottau K et al. 2020 SARS-CoV-2 in fruit bats, ferrets, pigs, and chickens: an experimental transmission study. Lancet Microbe 1, e218–e225. (doi:10.1016/S2666-5247(20)30089-6)

30. Babbitt GA, Fokoue EP, Srivastava HR, Callahan B, Rajendran M. 2022 Statistical machine learning for comparative protein dynamics with the DROIDS/maxDemon software pipeline. STAR Protoc. 3, 101194. (doi:10.1016/j.xpro.2022.101194)

31. Babbitt GA, Fokoue EP, Evans JR, Diller KI, Adams LE. 2020 DROIDS 3.0—Detecting Genetic and Drug Class Variant Impact on Conserved Protein Binding Dynamics. Biophys. J. 118, 541–551. (doi:10.1016/j.bpj.2019.12.008)

32. Babbitt GA, Mortensen JS, Coppola EE, Adams LE, Liao JK. 2018 DROIDS 1.20: A GUI-Based Pipeline for GPU-Accelerated Comparative Protein Dynamics. Biophys. J. 114, 1009–1017. (doi:10.1016/j.bpj.2018.01.020)

33. Rynkiewicz P, Lynch ML, Cui F, Hudson AO, Babbitt GA. 2021 Functional binding dynamics relevant to the evolution of zoonotic spillovers in endemic and emergent Betacoronavirus strains. J. Biomol. Struct. Dyn., 1–19. (doi:10.1080/07391102.2021.1953604)

34. Rajendran M, Ferran MC, Babbitt GA. 2022 Identifying vaccine escape sites via statistical comparisons of short-term molecular dynamics. Biophys. Rep. 0. (doi:10.1016/j.bpr.2022.100056)

35. Woods Group. 2005 GLYCAM Web. University of Georgia, Athens, GA: Complex Carbohydrate Research Center. See http://legacy.glycam.org.

36. Case DA et al. 2005 The Amber biomolecular simulation programs. J. Comput. Chem. 26, 1668–1688. (doi:10.1002/jcc.20290)

37. Pettersen EF, Goddard TD, Huang CC, Couch GS, Greenblatt DM, Meng EC, Ferrin TE. 2004 UCSF Chimera--a visualization system for exploratory research and analysis. J. Comput. Chem. 25, 1605–1612. (doi:10.1002/jcc.20084)

38. Fiser A, Do RK, Sali A. 2000 Modeling of loops in protein structures. Protein Sci. Publ. Protein Soc. 9, 1753–1773. (doi:10.1110/ps.9.9.1753)

39. Sali A, Blundell TL. 1993 Comparative protein modelling by satisfaction of spatial restraints. J. Mol. Biol. 234, 779–815. (doi:10.1006/jmbi.1993.1626)

40. Kirschner KN, Yongye AB, Tschampel SM, González-Outeiriño J, Daniels CR, Foley BL, Woods RJ. 2008 GLYCAM06: a generalizable biomolecular force field. Carbohydrates. J. Comput. Chem. 29, 622–655. (doi:10.1002/jcc.20820)

41. Babbitt GA, Lynch ML, McCoy M, Fokoue EP, Hudson AO. 2022 Function and evolution of B-Raf loop dynamics relevant to cancer recurrence under drug inhibition. J. Biomol. Struct. Dyn. 40, 468–483. (doi:10.1080/07391102.2020.1815578)

42. Ewald PP. 1921 Die Berechnung optischer und elektrostatischer Gitterpotentiale. Ann. Phys. 369, 253–287. (doi:10.1002/andp.19213690304)

43. Pierce LCT, Salomon-Ferrer R, Augusto F. de Oliveira C, McCammon JA, Walker RC. 2012 Routine Access to Millisecond Time Scale Events with Accelerated Molecular Dynamics. J. Chem. Theory Comput. 8, 2997–3002. (doi:10.1021/ct300284c)

44. Salomon-Ferrer R, Götz AW, Poole D, Le Grand S, Walker RC. 2013 Routine Microsecond Molecular Dynamics Simulations with AMBER on GPUs. 2. Explicit Solvent Particle Mesh Ewald. J. Chem. Theory Comput. 9, 3878–3888. (doi:10.1021/ct400314y)

45. Maier JA, Martinez C, Kasavajhala K, Wickstrom L, Hauser KE, Simmerling C. 2015 ff14SB: Improving the Accuracy of Protein Side Chain and Backbone Parameters from ff99SB. J. Chem. Theory Comput. 11, 3696–3713. (doi:10.1021/acs.jctc.5b00255)

46. Jorgensen WL, Chandrasekhar J, Madura JD, Impey RW, Klein ML. 1983 Comparison of simple potential functions for simulating liquid water. J. Chem. Phys. 79, 926–935. (doi:10.1063/1.445869)

47. Andersen HC. 1980 Molecular dynamics simulations at constant pressure and/or temperature. J. Chem. Phys. 72, 2384–2393. (doi:10.1063/1.439486)

48. Roe DR, Cheatham TE. 2013 PTRAJ and CPPTRAJ: Software for Processing and Analysis of Molecular Dynamics Trajectory Data. J. Chem. Theory Comput. 9, 3084–3095. (doi:10.1021/ct400341p)

49. Jumper J et al. 2021 Highly accurate protein structure prediction with AlphaFold. Nature 596, 583–589. (doi:10.1038/s41586-021-03819-2)

50. Hornak V, Abel R, Okur A, Strockbine B, Roitberg A, Simmerling C. 2006 Comparison of multiple Amber force fields and development of improved protein backbone parameters. Proteins 65, 712–725. (doi:10.1002/prot.21123)

51. Rehman Z, Fahim A, Bhatti MF. In press. Scouting the receptor-binding domain of SARS coronavirus 2: a comprehensive immunoinformatics inquisition. Future Virol., 10.2217/fvl-2020–0269. (doi:10.2217/fvl-2020-0269)

52. Liu K et al. 2021 Binding and molecular basis of the bat coronavirus RaTG13 virus to ACE2 in humans and other species. Cell 184, 3438–3451.e10. (doi:10.1016/j.cell.2021.05.031)

53. Heymann DL, Shindo N. 2020 COVID-19: what is next for public health? The Lancet 395, 542–545. (doi:10.1016/S0140-6736(20)30374-3)

54. Rothan HA, Byrareddy SN. 2020 The epidemiology and pathogenesis of coronavirus disease (COVID-19) outbreak. J. Autoimmun. 109, 102433. (doi:10.1016/j.jaut.2020.102433)

55. Wan Y, Shang J, Graham R, Baric RS, Li F. In press. Receptor Recognition by the Novel Coronavirus from Wuhan: an Analysis Based on Decade-Long Structural Studies of SARS Coronavirus. J. Virol. 94, e00127–20. (doi:10.1128/JVI.00127-20)

56. Munster VJ et al. 2020 Respiratory disease in rhesus macaques inoculated with SARS-CoV-2. Nature 585, 268–272. (doi:10.1038/s41586-020-2324-7)

57. Shi J et al. 2020 Susceptibility of ferrets, cats, dogs, and other domesticated animals to SARS-coronavirus 2. Science 368, 1016–1020. (doi:10.1126/science.abb7015)

58. Wrobel AG, Benton DJ, Xu P, Roustan C, Martin SR, Rosenthal PB, Skehel JJ, Gamblin SJ. 2020 SARS-CoV-2 and bat RaTG13 spike glycoprotein structures inform on virus evolution and furin-cleavage effects. Nat. Struct. Mol. Biol. 27, 763–767. (doi:10.1038/s41594-020-0468-7)

59. Neri A, Aygen M, Zukerman Z, Bahary C. 1980 Subjective assessment of sexual dysfunction of patients on long-term administration of digoxin. Arch. Sex. Behav. 9, 343–347. (doi:10.1007/BF01541359)

60. Seligson U, Söderlund C. 1986 ERCP and serum alkaline phosphatase in pancreatic carcinoma. Acta Chir. Scand. 152, 309–312.

61. Shivaprakash KN, Sen S, Paul S, Kiesecker JM, Bawa KS. 2021 Mammals, wildlife trade, and the next global pandemic. Curr. Biol. CB 31, 3671–3677.e3. (doi:10.1016/j.cub.2021.06.006)

62. de Morais HA, dos Santos AP, do Nascimento NC, Kmetiuk LB, Barbosa DS, Brandão PE, Guimarães AMS, Pettan-Brewer C, Biondo AW. 2020 Natural Infection by SARS-CoV-2 in Companion Animals: A Review of Case Reports and Current Evidence of Their Role in the Epidemiology of COVID-19. Front. Vet. Sci. 7.

63. Michelitsch A, Hoffmann D, Wernike K, Beer M. 2020 Occurrence of Antibodies against SARS-CoV-2 in the Domestic Cat Population of Germany. Vaccines 8, E772. (doi:10.3390/vaccines8040772)

64. Barrs VR et al. 2020 SARS-CoV-2 in Quarantined Domestic Cats from COVID-19 Households or Close Contacts, Hong Kong, China. Emerg. Infect. Dis. 26, 3071–3074. (doi:10.3201/eid2612.202786)

65. Patterson EI et al. 2020 Evidence of exposure to SARS-CoV-2 in cats and dogs from households in Italy. Nat. Commun. 11, 6231. (doi:10.1038/s41467-020-20097-0)

66. Fritz M et al. 2021 High prevalence of SARS-CoV-2 antibodies in pets from COVID-19+ households. One Health Amst. Neth. 11, 100192. (doi:10.1016/j.onehlt.2020.100192)

67. Domańska-Blicharz K et al. 2021 Mink SARS-CoV-2 Infection in Poland - Short Communication. J. Vet. Res. 65, 1–5. (doi:10.2478/jvetres-2021-0017)

68. Fenollar F, Mediannikov O, Maurin M, Devaux C, Colson P, Levasseur A, Fournier P-E, Raoult D. 2021 Mink, SARS-CoV-2, and the Human-Animal Interface. Front. Microbiol. 12, 663815. (doi:10.3389/fmicb.2021.663815)

