## Supplemental Figures for "Persistent cross-species SARS-CoV-2 variant infectivity predicted via comparative molecular dynamics simulation"

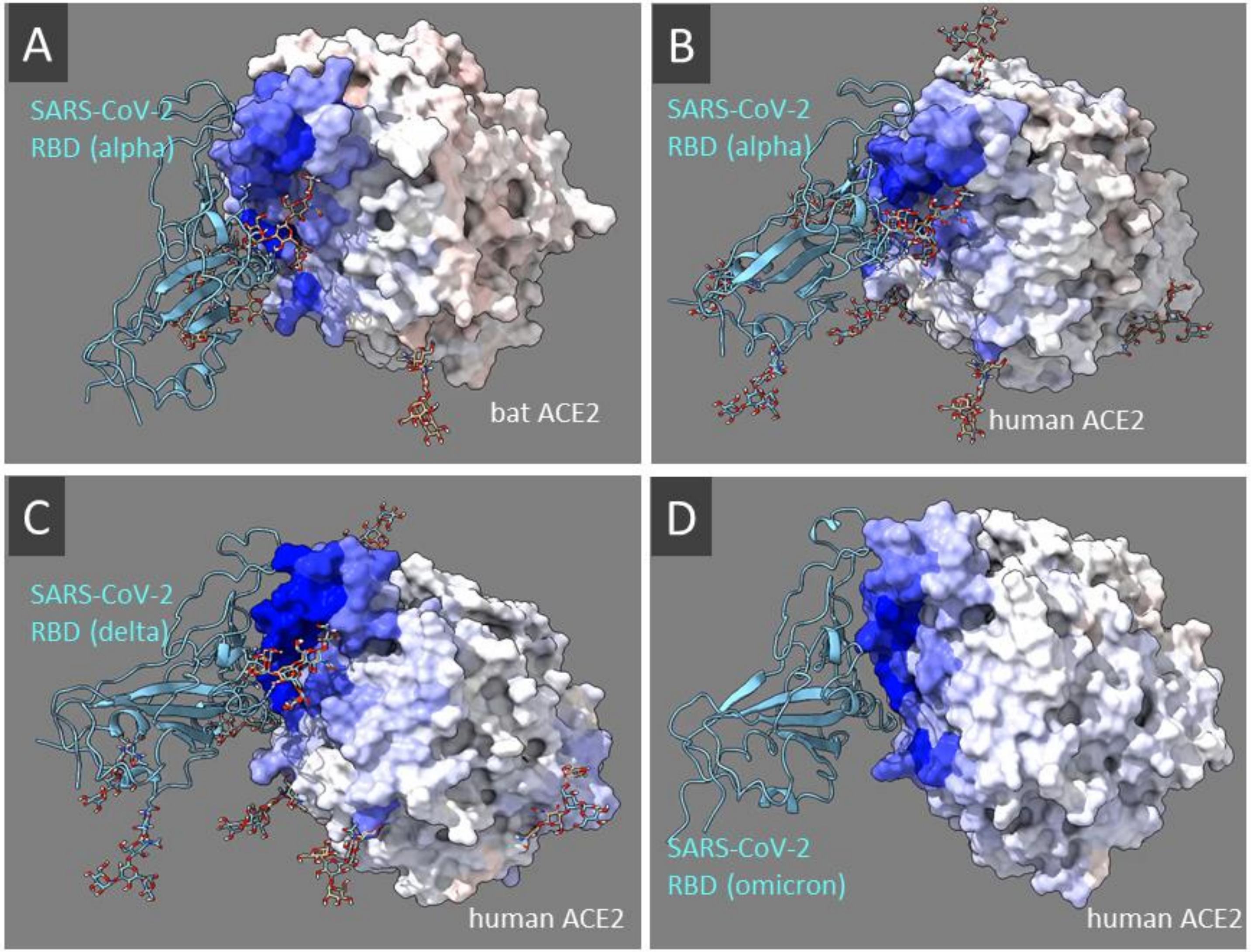


**Supplemental Figure 1. The binding signatures of SARS-CoV-2 receptor binding domain respective to human and bat (Rhinolophus macrotis) ACE2 orthologs.** The blue mapping indicates dampened atom fluctuation calculated as the signed KL divergence between site-wise distributions of atom fluctuation in the viral-bound vs unbound state of ACE2. The binding signatures for (A) reverse spillover of the SARS-CoV-2 alpha strain to bat ACE2 is compared to binding of (B) alpha, (C) delta, and (D) omicron strains to human ACE2.

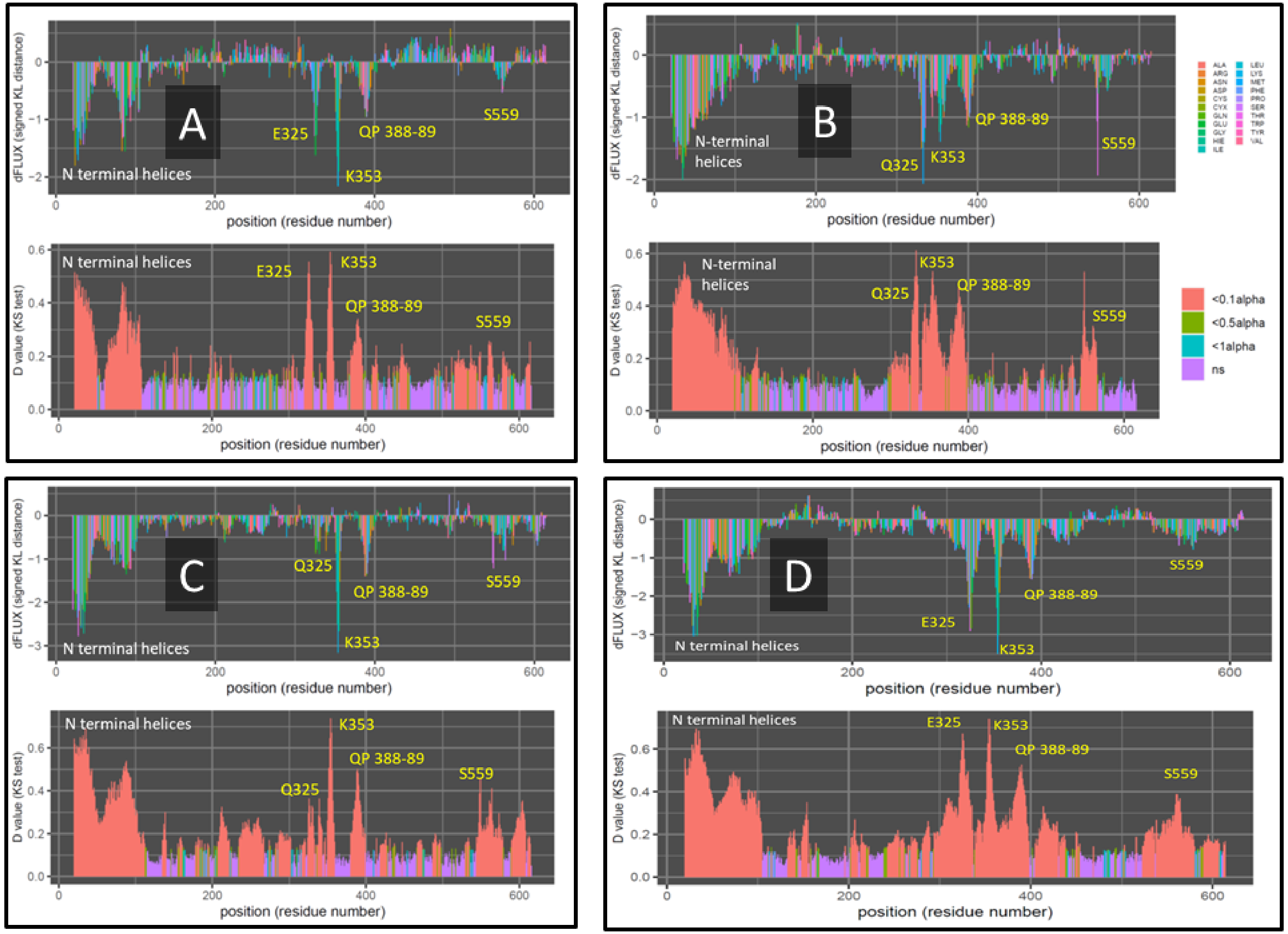


**Supplemental Figure 2. The binding signatures of SARS-CoV-2 receptor binding domain respective to human and bat (Rhinolophus macrotis) ACE2 orthologs.** The top plot indicates dampened atom fluctuation calculated as the signed KL divergence between site-wise distributions of atom fluctuation in the viral-bound vs unbound state of ACE2. The bottom plot indicates where these differences in atom fluctuation are significantly different according to Benjamini-Hochberg correscted two-sample Kolmogorov-Smirnov tests. The binding signatures for (A) reverse spillover of the SARS-CoV-2 alpha strain to bat ACE2 is compared to binding of (B) alpha, (C) delta, and (D) omicron strains to human ACE2.
